# Is galactose a hormetic sugar? Evidence from rat hippocampal redox regulatory network

**DOI:** 10.1101/2021.03.08.434370

**Authors:** J Homolak, Perhoc A Babic, A Knezovic, I Kodvanj, D Virag, Barilar J Osmanovic, P Riederer, M Salkovic-Petrisic

## Abstract

Galactose is a ubiquitous simple monosaccharide with yet incompletely understood biochemical and physiological role. Most of what we currently know about galactose is based on induction from the research on inherited disorders of galactose metabolism and animal models that exploit galactose-induced oxidative stress to model aging in rodents, however, recent evidence also demonstrates unique properties of galactose to conserve cellular function during the periods of starvation, and prevent and alleviate cognitive deficits in a rat model of sporadic Alzheimer’s disease. Here, we try to understand the molecular background of both detrimental and beneficial effects of galactose by exploring the acute systemic and hippocampal biochemical changes upon oral administration of galactose solution focusing primarily on the components of the redox regulatory network (RRN). Although orogastric gavage of galactose solution (200 mg/kg) was insufficient to induce systemic RRN disbalance in the first two hours upon administration, analysis of hippocampal RRN revealed a mild pro-oxidative shift accompanied by a paradoxical increase in tissue reductive capacity, suggesting overcompensation of endogenous antioxidant systems in the response to the pro-oxidative stimulus. The more thorough analysis revealed that galactose-induced increment of reductive capacity was accompanied by inflation of the hippocampal pool of nicotinamide adenine dinucleotide phosphates indicating ROS detoxification through disinhibition of the oxidative pentose phosphate pathway flux, reduced neuronal activity, and upregulation of Leloir pathway gatekeeper enzyme galactokinase-1. Based on the observed findings, and in the context of previous work on galactose, we propose a hormetic hypothesis of galactose action suggesting that the protective effects of galactose might be inseparable from its pro-oxidative effects at the biochemical level.

## Introduction

Galactose is a simple monosaccharide that was first discovered by Pasteur in 1856^1^. In structural terms, galactose differs from glucose, the most important energy substrate, only in the conFiguration of the hydroxyl group attached to the C4 carbon^2^. However, a plethora of evidence suggests that this modest structural difference enabled galactose to be selected by evolution to play an important biological role in a wide variety of living organisms. The importance of galactose is best illustrated in the following findings. Galactose, both in its free monosaccharide form, and attached to other molecules forming oligosaccharides, polysaccharides, glycoproteins, or glycolipids, has been identified in many living organisms including bacteria, plants, and animals^2^. Furthermore, the milk sugar lactose, known to be an essential nutrient for the development of nurslings, consists of galactose and glucose, and in rare lactose-free mammalian milk, found only in sea lions and marsupials, galactose is still present, usually in a trisaccharide form to provide energy and structural support for the developing brain^3^. Although the exact explanation why galactose has been selected by evolution as a component of lactose remains one of the biggest and most puzzling questions in biology^4^, its unique biochemical role offers some hints. Recent advancements in the field of glycobiology offer plenty of evidence that glycosylation serves a critical role in neural development, differentiation, axonal pathfinding, and synapses, explaining why a correct glycosylation pattern is a requirement for a functional nervous system^5, 6^. Galactose can be utilized in glycosylation through multiple biochemical pathways ^7^ and unlike glucose or mannose, the presence of galactose enables cells to maintain mature glycosylation patterns during times of sugar starvation^4^. In this context, current evidence suggests that the importance of galactose for living organisms might be related to its unique metabolic fate enabling galactose to ensure constant availability of glycosylation substrates. At the biochemical level, such an effect could be exerted by escaping strong regulatory mechanisms that are diverting metabolism exclusively towards energy generating systems during times of deficient nutrient availability. Interestingly, more than 160 years after its discovery, the physiological significance of galactose remains largely unknown, with the largest fraction of the accumulated scientific knowledge on galactose provided from two main sources: studies focused on galactosemia, a group of inherited disorders characterized by a reduced capacity of enzymes involved in galactose metabolism, and animal experiments focused on aging in which chronic parenteral administration of galactose to healthy animals is used to model oxidative stress-related pathophysiological mechanisms driving senescence. Both, however, offer limited understanding of the physiological significance of galactose pathways and whether they can be used to supply additional energy^8^, restore glycosylation patterns^4^, reduce ammonia load and promote cerebral amino acid synthesis^9^ without inducing oxidative stress with well known detrimental consequences^10^.

Galactosemia is sometimes referred to as proof of detrimental effects of galactose, however, a substantial body of evidence accumulated over the last few decades clearly shows that pathophysiological repercussions are a result of biochemical rerouting rather than the accumulation of the galactose substrate *per se*^11^. This is further corroborated by the fact that regardless of the specific enzyme deficiency, galactose accumulation occurs in all subtypes of galactosemia, but clinical manifestations differ greatly^11^. For example, galactokinase (**GALK; EC 2.7.1.6**) deficiency (Type II Galactosemia; MIM 230200) has been associated with a mild clinical presentation with cataracts often being the only evident abnormality^12^. In contrast, mutations in galactose-1-phosphate uridylyltransferase (**GALT; EC 2.7.7.12**) (Type I Galactosemia; MIM 230400), the second enzyme of the Leloir pathway, have been associated with potentially lethal consequences^13^. The exact pathomechanism responsible for these differences remains to be fully elucidated; however, accumulated evidence suggests that the dysfunction of this essential metabolic pathway, rather than the presence of galactose, is a mediator of the phenotypic expression of the disorder. For example, the build-up of galactose, galactitol, and galactonate has been reported in both GALK and GALT deficiency; however, patients suffering from Type I Galactosemia also accumulate galactose-1-phosphate (GAL-1-P) making it a potential molecular suspect responsible for the development of the pathological phenotype^11^. Even though the mechanism of toxicity of GAL-1-P remains a topic of debate, some authors suggest that competition with glucose-1-phosphate (GLC-1-P) for UTP-glucose-1-phosphate uridylyltransferase (**UGP; EC 2.7.7.9**) might lead to reduced availability of UDP-glucose substrates for UDP-glucose 4-epimerase (**GALE; EC 5.1.3.2**) further reducing UDP-galactose formation by impairment of a compensatory mechanism preserving regulatory lipid and protein glycosylation patterns in GALT-deficient cells^14^. Clinical findings on low-dose galactose supplementation in children with Type I Galactosemia further illustrate the complex pathophysiology underlying the disruption of galactose metabolism, as diet liberalization paradoxically restores N-glycosylation of IgG in a subset of patients, providing the theoretical groundwork for clinical improvement that remains to be confirmed^15^.

Another significant field of research that generated a considerable amount of information on the effects of galactose is the study of senescence where chronic parenteral galactose treatment has been used for decades for induction of oxidative stress-associated organismic degeneration. Interestingly, although a substantial number of studies take advantage of the model for the research of anti-aging therapeutics^16^, research focused on the elucidation of the pathophysiological background of galactose-induced senescence is scarce. Oxidative stress has been proposed as the main mechanism responsible for galactose-induced aging in rodents^16^, however, the exact mechanism has never been clarified. Two recent comprehensive reviews on galactose-induced aging^17^ and neurodegeneration^18^ propose three main mechanisms responsible for the pathophysiological effects of parenteral galactose. One potential explanation of galactose-induced oxidative stress suggests galactose metabolism generates a substantial amount of the alcohol galactitol through a reaction catalyzed by aldehyde reductase (**AR; EC 1.1.1.21**) with consequent depletion of the reduced form of nicotinamide adenine dinucleotide phosphate (NADPH) and osmotic stress, as accumulated galactitol cannot be further metabolized^17–20^. Moreover, it has been proposed that generation of Amadori products and Schiff bases by the reaction of galactose with free amino acid groups of proteins, lipids, and nucleic acid with consequent rearrangements producing advanced glycation end products (AGE) plays an important pathophysiological role^17, 18^. The third important mechanism suggests galactose generates H_2_O_2_ by undergoing an oxidation reaction catalyzed by galactose oxidase (**GO; EC 1.1.3.9**)^17, 18^ and some researchers suggest that this is, in fact, the only metabolic route of galactose that produces reactive oxygen species (ROS)^21^. Interestingly, although a substantial number of researchers working with galactose-induced models of aging in rodents refer to GO-mediated generation of H_2_O_2_ as an important mechanism explaining the detrimental effects of galactose ^17, 18, 22–34^, this is a fungal enzyme that doesn’t exist in rodents which makes these strong claims slightly suspicious as addressed in our recent comment^35^.

At the moment, questions of dose- and time-response effects, as well as differences in regards to the route of galactose administration and region-dependent metabolic capacities remain largely unaddressed, even though their implications are obvious and essential for understanding both the physiological and the pathophysiological significance of galactose treatment-related findings in the literature. In contrast to cognitive deterioration induced by chronic parenteral galactose administration to healthy rodents^36, 37^, we have found that galactose dissolved in drinking water (200 mg/kg/day) available ad libitum, can both prevent ^8^ and rescue ^38^ cognitive deficits in a rat model of sporadic Alzheimer’s disease induced by intracerebroventricular administration of streptozotocin (STZ-icv). Chogtu et al. reported both oral and subcutaneous galactose beneficially affect memory and learning; however, the observed effect was dependent on the duration of the treatment^39^. On the other hand, Budni et al. reported that even oral galactose treatment has the potential to induce cognitive impairment and oxidative damage in rats when given by oral gavage in a bolus dose once per day at relatively low doses (100 mg/kg)^40^.

Our research aimed to analyze the acute effects of a single dose of galactose administered by oral gavage. As oxidative stress is considered to be the main mediator of pathophysiological effects, we focused on the redox regulatory network (RRN), a set of intertwined subcellular systems responsible for the regulation of oxidant-antioxidant balance. We were especially interested in the cellular response to galactose-induced oxidative stress because oxidative stress has been recognized as an important mediator of hormesis^41^ - a dose-response phenomenon where low-dose stimulation with tolerable noxious stimulus provides long-term protection through the upregulation of protective cellular machinery. We hypothesized that acute responses of systemic RRN reflected by plasma mediators would help us understand whether even short-term increases of systemic galactose availability could induce redox disbalance before adaptive responses are activated. Moreover, we were interested in the response of hippocampal RRN, as the hippocampus (HPC) is considered to be one of the biological targets responsible for parenteral galactose-induced cognitive deficits.

## Materials and Methods

### Animals

Three-month-old male Wistar rats with a mean weight of 372 g (SD: 30) from the animal facility at the Department of Pharmacology (University of Zagreb School of Medicine) were used in the experiment. Animals were housed 2-3 per cage with a 12h light/dark cycle (7 am/7 pm). Room temperature and humidity were in the range of 21-23°C and 40-70%, respectively. Food pellets and water were available *ad libitum*. Standard wood-chip bedding was changed twice per week.

### Galactose treatment and experimental design

Animals were assigned to control (CTR) or one of galactose treatment groups (GAL 0.5h, GAL 1h, or GAL 2h) in a randomized order (8 animals/group). CTR received no galactose treatment and was used as baseline control and groups GAL 0.5h, GAL 1h, and GAL 2h received galactose solution by orogastric gavage (1 ml of 200 mg/kg galactose dissolved in water) and were sacrificed either 30, 60, or 120 minutes after the treatment respectively. Treatment dose was chosen based on our previous research^8, 38, 42^. Time-points were chosen based on previous findings that galactose plasma concentration is increased 15 minutes after both peroral and intraperitoneal treatment in both STZ-icv^38^ and control animals [unpublished data], and that this increment is resolved 30 minutes after peroral administration of 200 mg/kg solution (see Fig 2B) indicating that at approximately this time point galactose metabolism in target tissues is taking place.

### Tissue collection and preparation

Animals (6/group) were euthanized in general anesthesia (ketamine 50 mg/kg / xylazine 5 mg/kg ip) and decapitated. The brains were quickly removed and the hippocampi dissected, frozen in liquid nitrogen, and stored at −80°C. Afterward, the samples were subjected to three cycles of sonification (Microson Ultrasonic Cell Disruptor XL, Misonix, SAD) in five volumes of lysis buffer containing 10 mM HEPES, 1mM EDTA, 100 mM KCl, 1% Triton X-100, protease inhibitor cocktail (Sigma-Aldrich, USA) and phosphatase inhibitor (PhosSTOP, Roche, Switzerland) (pH 7.5) on ice. Homogenates were centrifuged for 10 minutes at 12000 RPM and 4°C. The protein concentration of supernatant for further analytical normalization was measured utilizing Lowry protein assay^43^ and supernatants were stored at −80°C until further analysis. The remaining animals (2/group) were subjected to deep anesthesia and transcardially perfused with 250 ml of saline followed by 250 ml of 4% paraformaldehyde (pH 7.4). Brains were quickly removed, cryoprotected through a series of sucrose (15%-30%), and stored at −80°C until further analysis. Cerebrospinal fluid (CSF) and plasma were extracted in deep anesthesia before decapitation or transcardial perfusion in all animals. CSF was sampled from the cisterna magna with a 330 μm insulin needle in fixed cervical flexion. Plasma was extracted from whole blood drawn from the retro-orbital sinus after centrifugation at 3600 RPM at 4°C for 10 minutes in heparinized tubes (100 μl per sample).

### Redox regulatory network analysis

To fully understand the effects of oral galactose-induced oxidative stress, we focused on redox regulatory network (RRN) subsystems. The superoxide detoxification subsystem (**Fig 1A**) was analyzed by 1,2,3-trihydroxybenzene autooxidation inhibition, the peroxidation subsystem (**Fig 1B**) was analyzed by [Co(CO_3_)_3_]Co-derived H_2_O_2_ dissociation rate and luminol chemiluminescence assay, lipid peroxidation (**Fig 1C**) was measured by TBARS, low molecular weight thiols (LMWT) (**Fig 1D**) were determined by TNB quantification after molecular weight fractionation, protein sulfhydryl content (**Fig 1E**) was calculated from the protein-DTNB reaction, and the overall reductive potential of the cell was examined by electrochemical determination of ORP and NRP/HistoNRP quantification (**Fig 1F**).

**Fig. 1.**
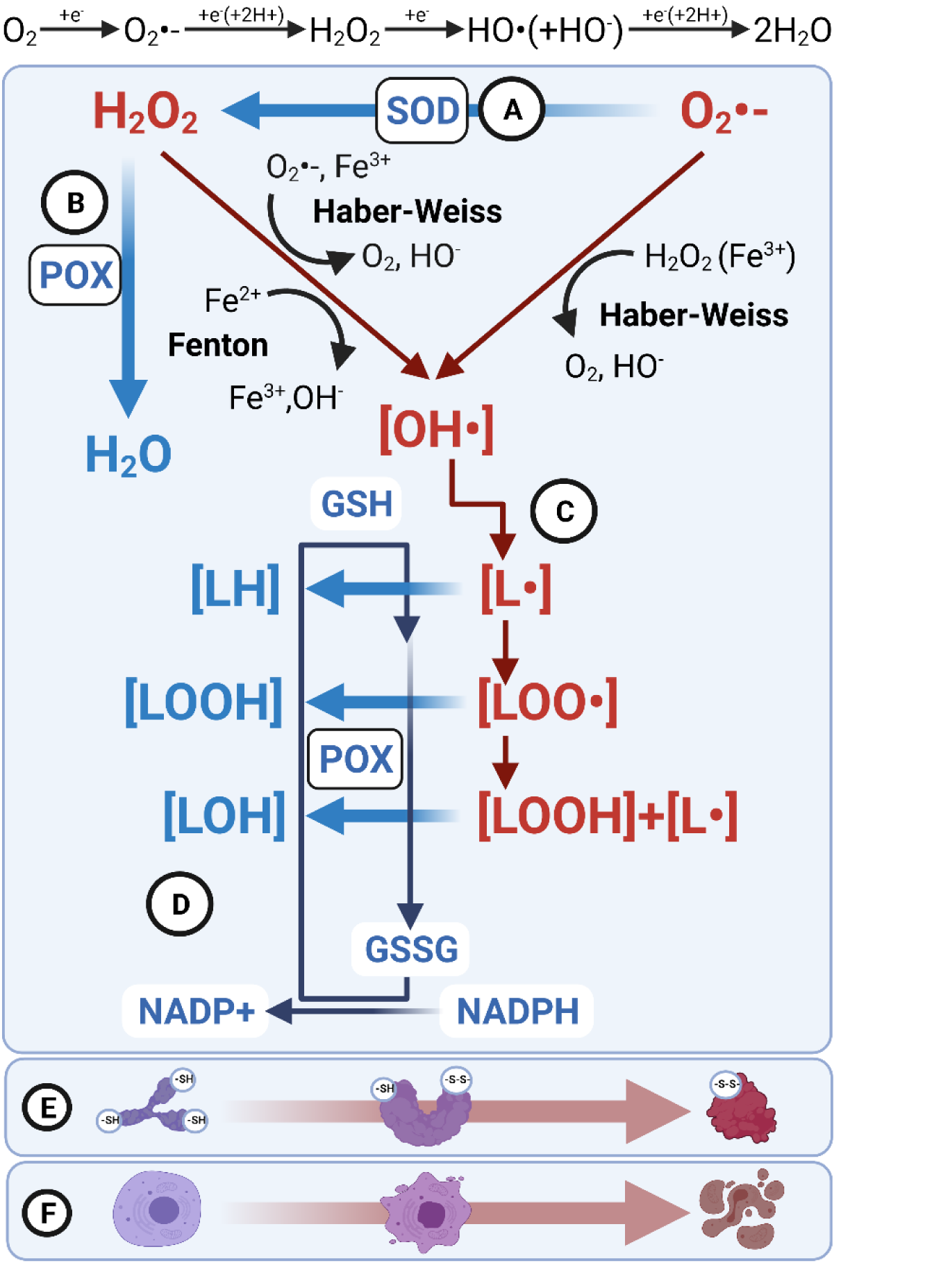
A simplified schematic overview of the most important cellular redox regulatory network subsystems explored in the study. For a deeper understanding of the chemical basis of reactive oxygen species (ROS), the reader is referred to the comprehensive and informative review of the chemical basis of ROS reactivity and the role in neurodegeneration by Fabrice Collin^44^. To maintain redox integrity, cell employs complementary systems that regulate the production and detoxification of important redox mediators. One of the most important sources of cellular ROS is the mitochondrial respiratory chain where 85% of O_2_ is metabolized during the successive 4 steps of 1-electron reduction reactions producing reactive intermediates^44^. Superoxide dismutase (SOD)**(A)** is an enzyme that catalyzes the dismutation of superoxide (O_2_•-) generated by the reduction of molecular oxygen. Dismutation of superoxide produces hydrogen peroxide (H_2_O_2_) an important endogenously produced reactive oxygen species (ROS) involved not only in ROS-mediated cellular damage but also in adaptor signaling. Cellular H_2_O_2_ is under the control of peroxidases **(B)**, a large group of enzymes that break down different cellular peroxides. One of the most important ubiquitously present peroxidases is catalase, an enzyme that catalyzes the decomposition of H_2_O_2_ to oxygen and water. Both H_2_O_2_ and O_2_•- have the potential to generate hydroxyl radicals (OH•) that in turn have the potential to generate lipid radicals (L•), lipid peroxy radicals (LOO•) and damaging lipid hydroperoxides (LOOH)**(C)**. Glutathione peroxidases are a major cellular defense system against the generation and propagation of lipid radicals. Glutathione peroxidase catalyzes both the reduction of H_2_O_2_ to water and the reduction of other peroxide radicals (e.g. lipid hydroperoxides) to alcohols and water. Glutathione peroxidases rely on cellular stores of monomeric glutathione (GSH), the most important low molecular cellular antioxidant that gets converted to a glutathione disulfide in the process (GSSG). Monomeric glutathione stores are then replenished by glutathione reductases that rely on the pool of reduced fraction of nicotinamide adenine dinucleotide phosphates NADPH to reduce glutathione disulfides **(D)**. Cellular proteins with abundant free sulfhydryl groups are involved in buffering excessive cellular ROS and it has been shown that this buffer pool has functional consequences **(E)**. For example, as briefly discussed in the text, glyceraldehyde 3-phosphate dehydrogenase, an important metabolic enzyme serves the role of pleiotropic molecular regulator integrating ROS sensing with cellular metabolism by forming intramolecular disulfides at active site cysteines, rendering the cell able to reroute substrates during the excessive ROS formation, and rapidly resume metabolic activity after oxidative stress subsides. On the other hand, excessive oxidative stress denatures proteins yielding an irreversible loss of function. Finally, at the cellular level, the harmonized function of the aforementioned redox-regulatory subsystems orchestrates the cellular fate when exposed to ROS-generating noxious stimulus. Low levels of oxidative stress that can be successfully compensated usually provoke adaptor response of the cell while excessive oxidative stress or moderate noxious stimulus in combination with a reduced capacity of one or several defense systems leads to the collapse of regulatory systems and results in cell death.

### Iodine/Potassium Iodide redox pair coupled oxidation-reduction potential

Iodine solutions (0.1 M I_2_ and 0.4 M KI) were used as the redox pair electrode solution in the experiment^45^. In short, 5 μL of hippocampal homogenate was mixed with 5 μL of the redox pair electrode solution and 40 μL of ddH2O. Samples were kept in the dark for 1 hour at room temperature. The reductive potential was measured with a 6230N Microprocessor meter (Jenco Instruments, San Diego, USA) coupled with a redox microsensor system ORP-146S (Shelf Scientific, Lazar Research Laboratories, USA). The system was composed of a Pt sensing element and Ag/AgCl reference with a fluoropolymer-based reference junction and capillary. KCl was used as a filling solution. The system was battery-powered during all measurements to exclude potential powerline interference. The oxidation-reduction potential measurement range was −1500 mV to +1500 mV with a system accuracy of ±0.5 mV.

### Hydrogen peroxide dissociation rate

Catalase and peroxidase activity was determined indirectly by hydrogen peroxide dissociation rate. In short, samples were vortexed and 30 μL of each sample was placed in a 96-well plate. Sample background absorbance was checked at 450 and 560 nm to control for internal interference. H_2_O_2_ concentration was determined by oxidation of cobalt (II) to cobalt (III) in the presence of bicarbonate ions by spectrophotometric quantification of carbonato-cobaltate (III) complex ([Co(CO_3_)_3_]Co) at a 450 nm peak. Reference absorbance for each sample was estimated by baseline H_2_O_2_ quantification at t_0_= 0 s, and final absorbance was determined at t_1_= 120 s. The reaction was started by adding 40 μL of H_2_O_2_ solution (10 mM in 1xPBS) and stopped by adding 100 μL of cobalt (II) hexametaphosphate working solution as previously described by Hadwan^46^. Absorbance was measured with Infinite F200 PRO multimodal microplate reader (Tecan, Switzerland).

### Superoxide dismutase activity

Superoxide dismutase activity was measured by 1,2,3-trihydroxybenzene autooxidation inhibition^47^. In short, 15 μL of 60 mM 1,2,3-trihydroxybenzene dissolved in 1 mM HCl was added to 1000 μL of 0.05 M Tris-HCl and 1mM Na_2_EDTA (pH 8.2), briefly vortexed and mixed with 10 μL of hippocampal homogenate. Absorbance increase was recorded at 325 nm for 300 seconds. Maximal 1,2,3-trihydroxybenzene autooxidation was determined by the same procedure without adding the tissue sample. Autooxidation inhibition was calculated as a ratio of sample and reference sample difference in end-point and baseline absorbance, ratiometrically corrected for tissue sample protein concentration. Continuous change in absorbance was determined by CamSpec M350 DoubleBeam UV-Visible Spectrophotometer (Cambridge, UK).

### Low molecular weight thiols and total protein sulfhydryl content

Low molecular weight thiols (LMWT) and tissue protein sulfhydryl content were assessed by the formation of 5-thio-2-nitrobenzoic acid (TNB) in a reaction between sulfhydryl groups and 5,5’-dithio-bis(2-nitrobenzoic acid) (DTNB)^48, 49^. In short, 25 μL of tissue homogenate was incubated with 25 μL of sulfosalicylic acid (4% w/v) for 1h on ice and centrifuged for 10 minutes at 10000 RPM. The supernatant (30 μL) was transferred to separate tubes for LMWT determination. The protein pellet was mixed with 30 μL of DTNB (4 mg/ml in 5% sodium citrate) and 540 μL KPB (0.1 M, pH 7.4), and the mixture absorbance was read at 412 nm for total protein sulfhydryl content determination. The remaining supernatant was added to the same DTNB reagent and KPB and its absorbance measured at 412 nm were used for LMWT estimation. Absorbance was determined by CamSpec M350 DoubleBeam UV-Visible Spectrophotometer (Cambridge, UK). The calculated sulfhydryl content was ratiometrically corrected for sample protein concentration.

### Lipid peroxidation

Lipid peroxidation was estimated employing standard malondialdehyde assay based on the determination of thiobarbituric acid reactive substances (TBARS)^48^. In short, 36 μL of tissue homogenate was mixed with 400 μL TBARS-TCA reagent (0,375% thiobarbituric acid; 15% trichloroacetic acid) and 164 μL of ddH_2_O. Samples were incubated at 95°C for 20 minutes in perforated Eppendorf tubes and left to cool down under tap water. The complex of thiobarbituric acid and malondialdehyde was extracted by adding 600 μL of n-butanol to the mixture. The absorbance of the butanol fraction was analyzed at 532 nm and the amount of TBARS was estimated based on the molar extinction coefficient of 1,56×10^5^ M^-1^ cm^-1^. Absorbance was determined by CamSpec M350 DoubleBeam UV-Visible Spectrophotometer (Cambridge, UK). Calculated TBARS content was corrected for sample protein concentration.

### Cytochrome c-mediated peroxide dissociation

Cytochrome c-mediated hydrogen peroxide dissociation was quantified by native sodium dodecyl sulfate-polyacrylamide gel electrophoresis (SDS-PAGE) followed by membrane chemiluminiscence^50^. The protocol was the same as described for western blot, with the omission of heating at 95°C during sample preparation. After electrophoretic separation and transfer, the nitrocellulose membrane was incubated in an enhanced chemiluminescence reagent (SuperSignal™ West Femto; Thermo Fisher Scientific, USA). Luminescence was captured by MicroChemi high-performance chemiluminescence imager (DNR Bio-Imaging Systems Ltd., Israel) and GelCapture software. The consequent analysis was the same as described for Western blot.

### Western blot

Protein expression of c-fos and GALK1 was measured by Western blot with subsequent chemiluminescent-based quantification. Equal amounts of total protein (35 μg) determined by the Lowry assay were separated by SDS-PAGE (9% polyacrylamide gels) and transferred to nitrocellulose membranes using the wet transfer (100V/400mA/60min in 140 mM glycine, 18 mM Tris, 20% methanol (v/v)). Nitrocellulose membranes were incubated in blocking buffer (5% nonfat dry milk solution with 0,5% Tween 20 in low-salt washing buffer (LSWB: 10 mM Tris, 150 mM NaCl, pH 7.5)) for 1h at room temperature. Blocked membranes were incubated in c-fos (1:000, Cell Signaling Technology, USA) or GALK1 (1:1000, Invitrogen, USA) primary antibody solution in blocking buffer overnight at 4°C. After incubation, membranes were washed 3x in LSWB and incubated with anti-rabbit IgG secondary antibody (1:2000, Cell Signaing Technology, USA) in blocking buffer. Membranes were washed 3x in LSWB and incubated in chemiluminescent reagent (SuperSignal™ West Femto; Thermo Fisher Scientific, USA). Luminescence was captured by MicroChemi high-performance chemiluminescence imager (DNR Bio-Imaging Systems Ltd., Israel) and GelCapture software. The same procedure was repeated for β-actin (1:2000, Sigma-Aldrich, USA) used as a loading control. Immunoblot densitometry was done in Fiji (NIH, USA) using Gel Analyzer protocol after Rolling Ball background subtraction.

### Nitrocellulose redox permanganometry

Nitrocellulose redox permanganometry (NRP)^51^ was used for reductive potential determination in plasma and hippocampal homogenates. HistoNRP was used for the reductive potential analysis of brain tissue cryosections. In short, plasma samples were centrifuged and 1 μL of each sample was placed onto a clean piece of nitrocellulose membrane (Amersham Protran 0.45; GE Healthcare Life Sciences, USA) and left to dry out. Once dry, the membrane was immersed in NRP reagent (0.2 g KMnO_4_ in 20 ml ddH_2_O) for 30 seconds. The reaction was terminated and MnO_2_ contrast was developed by placing the membrane under running ddH_2_O. For the HistoNRP experiment, we used 30 μm thick brain cryosections (Leica CM1850; Leica Biosystems, USA). In short, sections were mounted onto microscope slides and tissue proteins were transferred onto nitrocellulose by passive diffusion slice print blotting protocol with PBS as a transfer buffer. The transfer was conducted overnight under the constant pressure of 31.384 mmHg. Membranes were carefully removed with PBS, left to dry out, and reacted with NRP solution using the same protocol as described above. All membranes were digitalized by an office scanner (SP 4410SF, RICOH, USA) and analyzed in Fiji (NIH, USA) as described in detail in^51^.

### Total NADP and NADPH measurement

Total NADP and NADPH in hippocampal homogenates were measured colorimetrically using NADP/NADPH Quantitation kit (Sigma-Aldrich, USA) following the manufacturer’s instructions. Absorbance was read with Infinite F200 PRO multimodal microplate reader (Tecan, Switzerland).

### Glutamate quantification

Glutamate quantity in hippocampal homogenates (10 μl) was measured colorimetrically utilizing Glutamate Assay Kit (Sigma-Aldrich, USA) following the manufacturer’s instructions. Absorbance at 450 nm was read with Infinite F200 PRO multimodal microplate reader (Tecan, Switzerland).

### Plasma glucose and galactose measurement

Glucose was measured in 10 μl of plasma sample colorimetrically with the standard Trinder method utilizing glucose oxidase and 4-amino-antipyrine provided in a commercial kit (Greiner Diagnostic, Germany). Plasma galactose was determined with the Amplex™ Red Galactose/Galactose Oxidase Assay Kit (Thermo Fisher Scientific, USA) following the manufacturer’s instructions. Absorbance was read at 570 nm with Infinite F200 PRO multimodal microplate reader (Tecan, Switzerland).

### Data analysis and visualization

Data analysis and visualization were conducted in R (4.0.2). As our experiment was exploratory raw data was reported for all variables measured both in plasma and hippocampal homogenates. The association of variables of interest was explored by principal component analysis on scaled and mean-imputed datasets and reported as a biplot of individual animals. Individual variables and their contributions to the 1^st^ and 2^nd^ dimension were reported as vectors in the biplot space with color and length signaling contributions. Coordinates of individual animals in the biplot space were used for post-hoc group comparisons. The Wilcoxon signed-rank test was used as a non-parametric test for the comparison of two groups with an alpha set at 5%.

## Results

### The effect of acute oral galactose treatment on plasma redox regulatory network

To examine how acute oral galactose treatment affects overall systemic RRN, we first analyzed plasma samples following the assumption that widespread systemic oxidative stress should be mirrored by plasma redox status. Analysis of SOD activity revealed reduced activity 0.5h after galactose treatment, with normalization after 1h (**Fig 2A**). LMWT followed a similar trend, but group differences were not statistically significant (GAL 0.5h vs GAL 2h; p=0.13) (**Fig 2B**). Interestingly, in contrast to what we expected, lipid peroxidation end products were reduced both 0.5h and 1h after oral galactose treatment in comparison to controls (**Fig 2C**). Overall oxidation-reduction potential showed a trend of increment after galactose treatment with no statistical significance (CTR vs GAL; p=0.15) (**Fig 2D**). Another marker of reductive potential, NRP, was also unchanged with a slight trend of increment only in GAL 2h in comparison to GAL 1h (p=0.14) (**Fig 2E**). Finally, we measured plasma glucose to assess how single oral galactose treatment effects were related to glucose perturbations, with no significant changes being detected in analyzed time points (**Fig 2F**). Furthermore, no difference in plasma galactose concentration was observed, indicating the administered galactose was fully absorbed and/or metabolized by target organs in the first 30 minutes (**Fig 2G**). Overall, apart from reduced lipid peroxidation, none of the measured parameters indicated acute oral galactose treatment in the dose of 200mg/kg was able to perturb rat systemic redox regulatory network. Multivariate analysis of measured indicators revealed subtle clusters that described 31.2% and 25.5% of the variance in the 1^st^ and 2^nd^ dimensions respectively. Coordinates of individual animals in the biplot are shown in **Fig 2H**, and the contribution of individual variables is depicted in **Fig 2I.** Biplot coordinates of individual animals classified by treatment in respect to the first two principal components are shown in **Fig 2J**, and the overall effect of galactose in respect to the generated biplot is shown in **Fig 2K**. Subtle clusters suggested galactose treatment exerted a less time-dependent effect on the ORP, NRP, and LMWT and a more time-dependent effect on SOD activity and glucose concentration.

**Fig 2.**
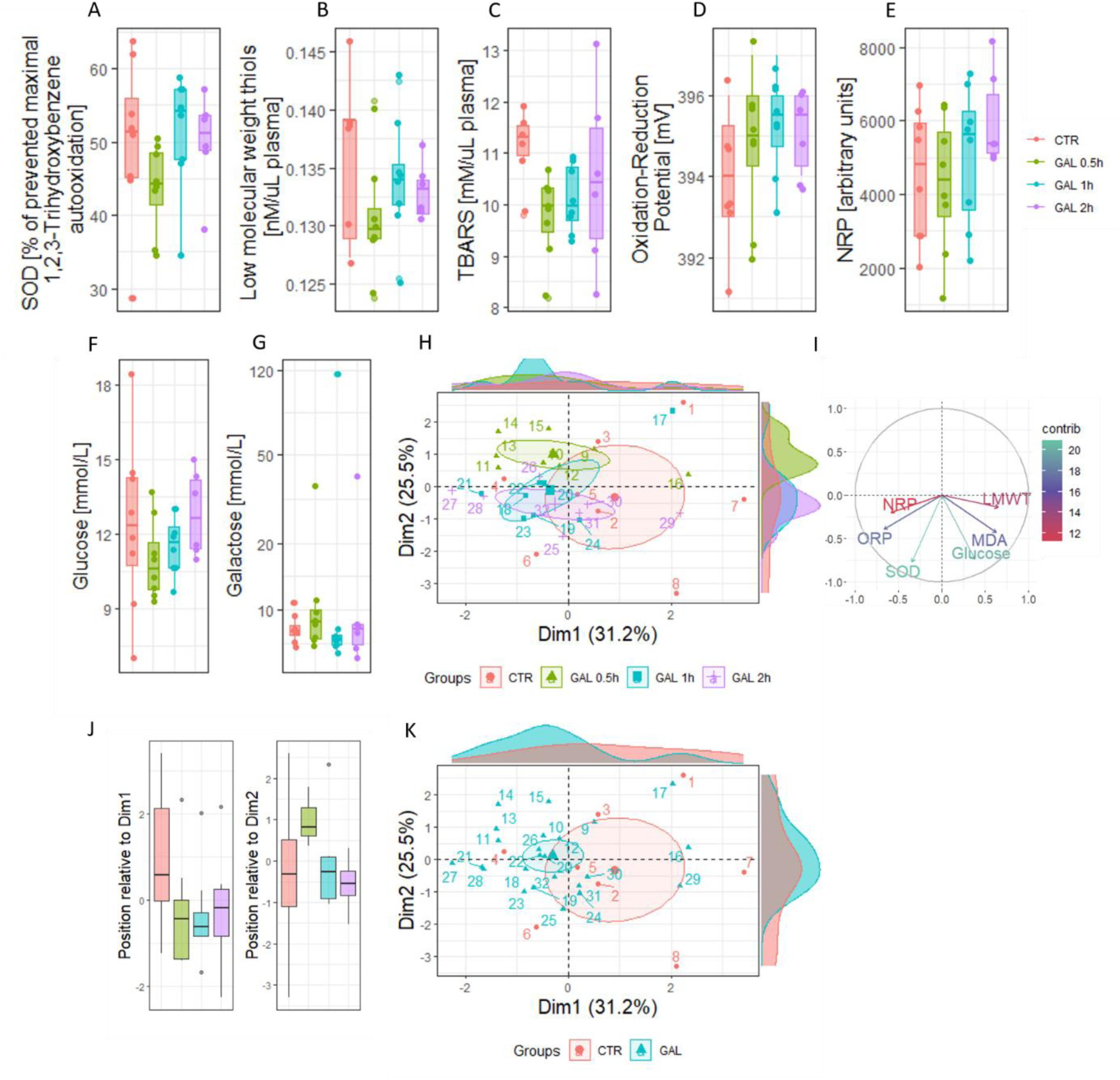
The effect of acute oral galactose treatment on the plasma redox regulatory network. The effect of acute oral galactose treatment on **(A)** the activity of superoxide dismutase (SOD)[CTR vs GAL 0.5h p=.065; GAL 0.g5 vs GAL 1h p=.052], **(B)** the concentration of low molecular weight thiols (LMWT), **(C)** lipid peroxidation reflected by the concentration of thiobarbituric acid reactive substances (TBARS)[CTR vs GAL 0.5h p=.017; CTR vs GAL 1h p=.023], overall redox balance measured by **(D)** I_2_/KI redox couple-stabilized oxidation-reduction potential (ORP), and **(E)** nitrocellulose redox permanganometry (NRP). **(F)** The effect of acute oral galactose treatment on plasma glucose concentration [GAL 0.5h vs GAL 2h p=.053]. **(G)** The effect of acute oral galactose treatment on plasma galactose concentration. **(H)** Principal component analysis biplot of the 1^st^ and 2^nd^ principal components with individual animal coordinates. **(I)** Contribution of individual variables to the 1^st^ and 2^nd^ principal components with a contribution (%) depicted by color and length of the vector. **(J)** Comparison of animals by groups based on assigned appropriate coordinates in respect to the 1^st^ and 2^nd^ component respectively. **(K)** The “overall” effect of galactose demonstrated on the biplot shown in Fig 2G. CTR – control; GAL – galactose treated. * [Wilcoxon signed-rank comparisons].

### The effect of acute oral galactose treatment on hippocampal redox regulatory network

We then moved on to examine RRN subsystems in the hippocampus. SOD activity was reduced in all groups after galactose treatment in comparison with controls (**Fig 3A**). Peroxidase activity approximated from the H_2_O_2_ dissociation rate demonstrated an increasing trend reaching the highest values in the GAL 1h group (**Fig 3B**). Here, we additionally tested cytochrome C H_2_O_2_ dissociation activity and observed a similar trend with the highest activity in the GAL 1h group [CTR vs GAL 1h; p=0.043] (**Fig 3C**). LMWT concentration remained unchanged in all groups (**Fig 3D**), but total protein-free sulfhydryl groups demonstrated a transient decrement upon treatment normalized over time [GAL 0.5h vs GAL 2h; p=0.037] (**Fig 3E**). Lipid peroxidation remained largely unchanged with a trend of increment observed only in GAL 1h in comparison to GAL 0.5h [p=0.12] (**Fig 3F**). We then moved on to examine the effect of oral galactose on the overall cellular redox balance. Based on the results obtained from the analysis of individual RRN subsystems (reduced SOD activity, increased peroxidase activities, possible transient reduction in total protein free sulfhydryl groups, and a trend towards increased lipid peroxidation) we expected to see the redox balance shifted towards a pro-oxidative state. Interestingly, in contrast to our expectations, ORP was reduced rather than increased in the GAL 1h group, suggesting galactose treatment increased reductive capacity (**Fig 3G**). Although ORP changes were consistent, the effect was small and we wanted to verify our findings; therefore, we performed an additional examination of the reductive potential using NRP, where the finding of an increased reduction potential after galactose treatment was confirmed (**Fig 3H**). However, as ORP and NRP suggested a different pattern of reductive potential increment, we analyzed their correlation by linear regression, where we obtained a negative correlation, as expected (R=-0.51; p=0.013). Interestingly, when treatment control was introduced to the model, the effect was preserved in the control group, but lost in galactose-treated rats (CTR: R=-0.88; p=0.02/GAL: R=-0.035; p=0.9) suggesting that galactose treatment interfered with a reductive component of the system that enters a redox reaction with KMnO_4_ much more effectively than with the I_2_/KI redox pair used for ORP stabilization, as described in the Methods section (**Fig 3I**). To further explore the nature of this unexpected rise in the total reductive potential induced by acute oral galactose treatment, we conducted a HistoNRP^51^ analysis on brain tissue cryosections. As shown in **Fig 4A** and **Fig 4B**, galactose induced an increase in overall brain reductive capacity in a time- and region-dependent manner, with the greatest values recorded in the GAL 1h group, and with values in the GAL 2h group being comparable to those recorded in the CTR group (or lower). As some differences in the regional distribution of the reductive potential increment were observed, a separate analysis of the specific brain areas of interest was conducted (**Fig 4C**). Even though most of the analyzed brain areas demonstrated a similar pattern with a steady increment up to the GAL 1h time-point and normalization or hyper compensation in the GAL 2h time-point, some areas, such as the dorsal hippocampus and parietal cortex, demonstrated a steady progressive galactose-induced decrement of the reductive capacity, indicating a variable redox or metabolic tolerance to galactose throughout the brain (**Fig 4A, Fig 4C**). Interestingly, a different pattern was observed for the ventral and dorsal portions of the hippocampus indicating that different subsets of cells inside the same anatomical region could differ regarding the galactose metabolic tolerance. Especially high increment of reductive capacity was observed in substantia nigra, as evident both in **Fig 4A** and **Fig 4C**.

**Fig 3.**
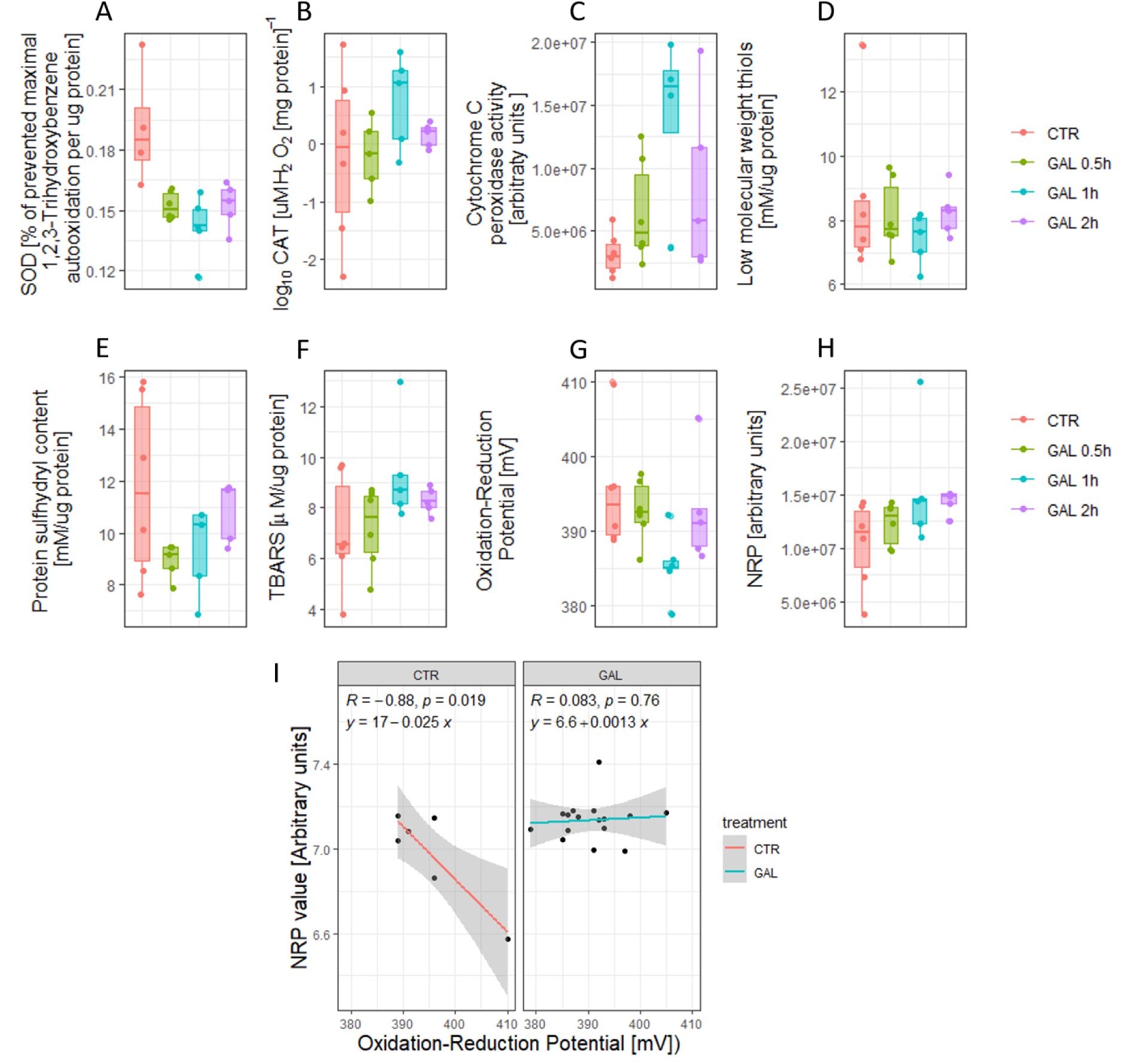
The effect of acute oral galactose treatment on the hippocampal redox regulatory network. The effect of acute oral galactose treatment on **(A)** the activity of superoxide dismutase (SOD)[CTR vs GAL 0.5h p=.009; vs GAL 1h p=.016; vs GAL 2h p=.032], **(B)** H_2_O_2_ dissociation rate, **(C)** peroxidase-like activity of cytochrome c [CTR vs GAL 1h p=.038], **(D)** low molecular weight thiols (LMWT), **(E)** protein free sulfhydryl groups [GAL 0.5hh vs GAL 2h p=.032], **(F)** lipid peroxidation reflected by the concentration of thiobarbituric acid reactive substances (TBARS), and the overall redox balance measured by **(G)** I_2_/KI redox couple-stabilized oxidation-reduction potential (ORP) [GAL 1h vs CTR p=.035; vs GAL 0.5h p=.035; vs GAL 2h p=.059], and **(H)** nitrocellulose redox permanganometry (NRP) [CTR vs GAL 2h p=.030; GAL 0.5h vs GAL 2h p=.052]. **(I)** Linear regression of NRP and ORP by treatment regardless of the time-point in the GAL group. CTR – control; GAL – galactose treated. * [Wilcoxon signed-rank comparisons].

We then moved on to examine the possible background of the observed effect. Here, we focused on the NADP system for several reasons. It has been shown that acute oxidative stress challenge exerts a disinhibitory effect on the evolutionary conserved protective pentose phosphate pathway (PPP) metabolic network^52–55^. Moreover, galactose can enter PPP at various stages, and its conversion to glucose-6-phosphate provides a biochemical substrate for the oxidative branch of PPP. Furthermore, previous findings in activated T cells grown on galactose-containing medium suggest galactose-derived glucose is shuttled into the PPP rather than the glycolytic pathway ^56, 57^ and recent evidence shows that galactose, in contrast to glucose, doesn’t activate the glycolytic gatekeeper enzyme phosphofructokinase-1 (PFK-1)^4^ (**Fig 5**). We first analyzed NADPH, a reductive equivalent produced in the pentose phosphate pathway and used for regeneration of cellular antioxidant systems (**Fig 1C**) and anabolic biosynthesis. As shown in **Fig 4D**, NADPH concentration remained unchanged with a slight trend of increment after GAL treatment [CTR vs GAL; p=0.14]. However, analysis of total NADP revealed galactose treatment increased the concentration of total NADP by 542% [CTR vs GAL; p=0.0017] (**Fig 4E**). Conversely, the ratio of reduced to total NADP fraction was decreased following galactose treatment (**Fig 4F**) indicating increased utilization of reductive equivalents, possibly for the replenishment of cellular antioxidant systems and providing a potential explanation for the paradoxical pro-oxidant shift of some RRN subsystems with a simultaneous increment of reductive capacity. We further hypothesized that the observed activation of the protective oxidative pentose phosphate shunt would be followed by a reduction of cellular activity to compensate for the reduced substrate flux into the lower glycolysis pathway and further decrease aerobic respiration-generated ROS. To test this, we analyzed the expression of c-fos, a marker of neuronal activity, and found its expression to be reduced by GAL treatment in a time-dependent manner (**Fig 4G**). Moreover, we expected that acute galactose treatment would induce the expression of galactokinase-1 (GALK1), an enzyme that catalyzes the first committed step of the Leloir pathway as an adaptive response to increased galactose availability. As shown in **Fig 4H**, galactose treatment enhanced the expression of GALK1 in a time-dependent manner, with the increase being most pronounced 2h after the treatment [CTR vs GAL 2h; +222%; p=0.055]. Furthermore, as it was previously suggested by Reutter and colleagues that one of the potential neuroprotective mechanisms of galactose might be mediated by its ability to consume equivalents of ammonia, we additionally tested the effect of galactose on the concentration of hippocampal glutamate and observed a rapid reduction with subsequent normalization after 2h (**Fig 4I**). The association between individual metabolic and redox-related parameters was further explored by principal component analysis reported as a biplot with coordinates of individual animals classified based on treatment group (**Fig 4J**) or the “overall effect of galactose treatment” (**Fig 4K**). Contribution of individual parameters to the 1^st^ and 2^nd^ dimension are shown in **Fig 4L**, and comparison of animals by groups based on assigned appropriate coordinates in **Fig 4M.**

**Fig 4.**
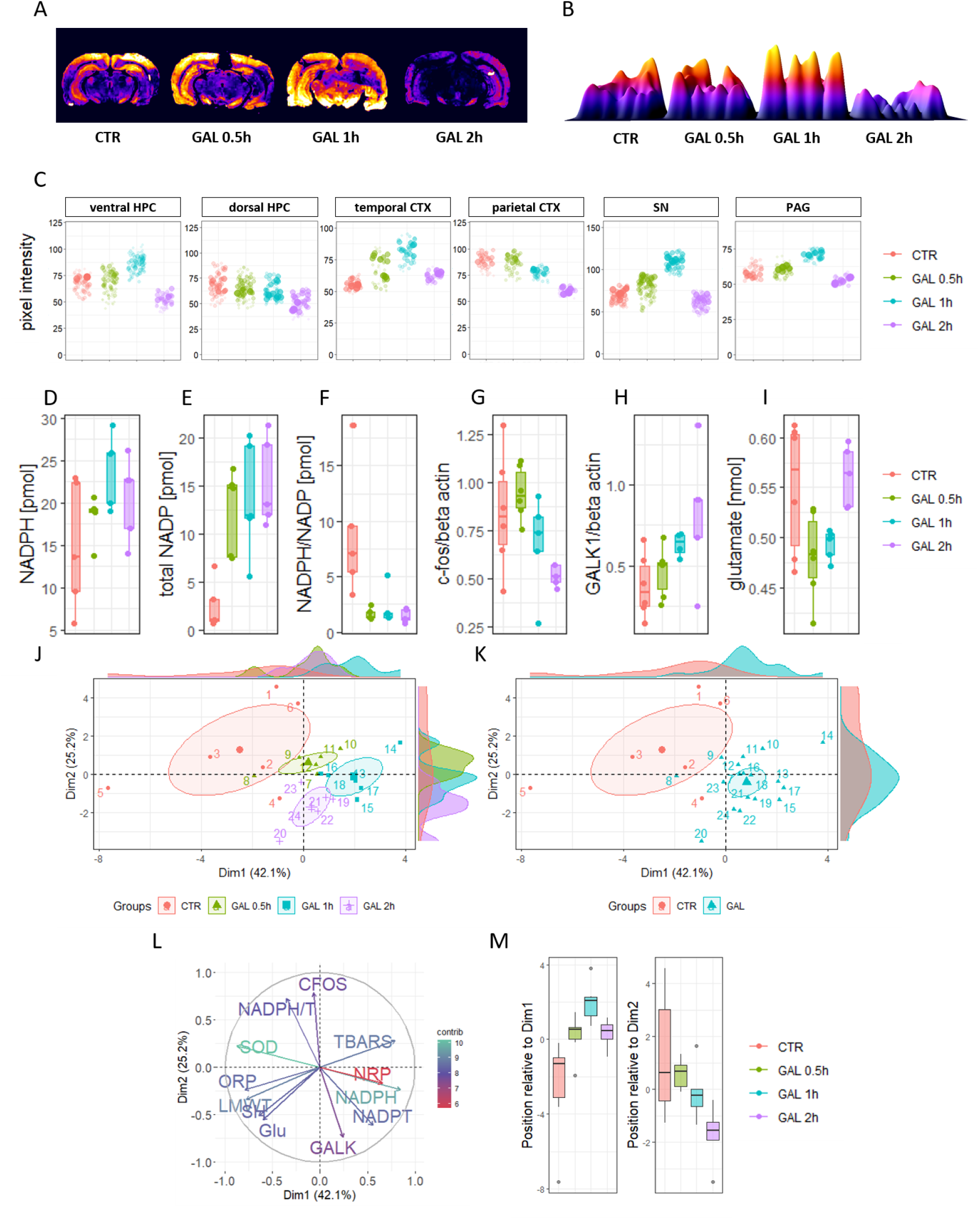
Reductive capacity, nicotinamide adenine dinucleotide phosphate (NADPH) system, and metabolic parameters in the rat brain following acute oral galactose treatment. Histo-nitrocellulose redox permanganometry (HistoNRP) of representative brain tissue sections **(A)** and 3D surface plots of the same animals **(B)**. **(C)** NRP-obtained values (greater value indicated increased reductive capacity) represented by individual pixel intensities from 6 different anatomical regions of the brain. **(D-M)** represent hippocampal samples. Acute oral galactose treatment effect on **(D)** hippocampal NADPH, **(E)** total NADP [CTR vs GAL 0.5h p=.007; vs GAL 1h p=.016; vs GAL 2h p=.007], **(F)** ratio of reduced to total NADP [CTR vs GAL 0.5h p=.007; vs GAL 1h p=.016; vs GAL 2h p=.007], **(G)** expression of c-fos [CTR vs GAL 2h p=.082; GAL 0.5h vs GAL 1h p=.052;], and **(H)** galactokinase-1 (GALK-1) normalized to beta-actin [CTR vs GAL 1h p=.038; GAL 0.5h vs GAL 1h p=.038], and concentration of **(I)** tissue glutamate [GAL 0.5h vs GAL 2h p=.004; GAL 1h vs GAL 2h p=.007]. Principal component analysis biplot of the 1^st^ and 2^nd^ principal components with individual animal coordinates based on the group **(J)** or “overall treatment with GAL” **(K)**. **(L)** Contribution of individual variables to the 1^st^ and 2^nd^ principal components with a contribution (%) depicted by color and length of the vector. **(M)** Comparison of animals by groups based on assigned appropriate coordinates in respect to the 1^st^ and 2^nd^ component respectively [Dim1: CTR vs GAL 0.5h p=.026; vs GAL 1h p=.002; vs GAL 2h p=.004. GAL 0.5h vs GAL 1h p=.008. GAL 1h vs GAL 2h p=.015. Dim2: CTR vs GAL 2h p=.015. GAL 1h vs GAL 2h p=.026.]. CTR – control; GAL – galactose; HPC – hippocampus; CTX – cortex: SN – substantia nigra; PAG – periaqueductal gray. * [Wilcoxon signed-rank comparisons].

**Fig 5.**
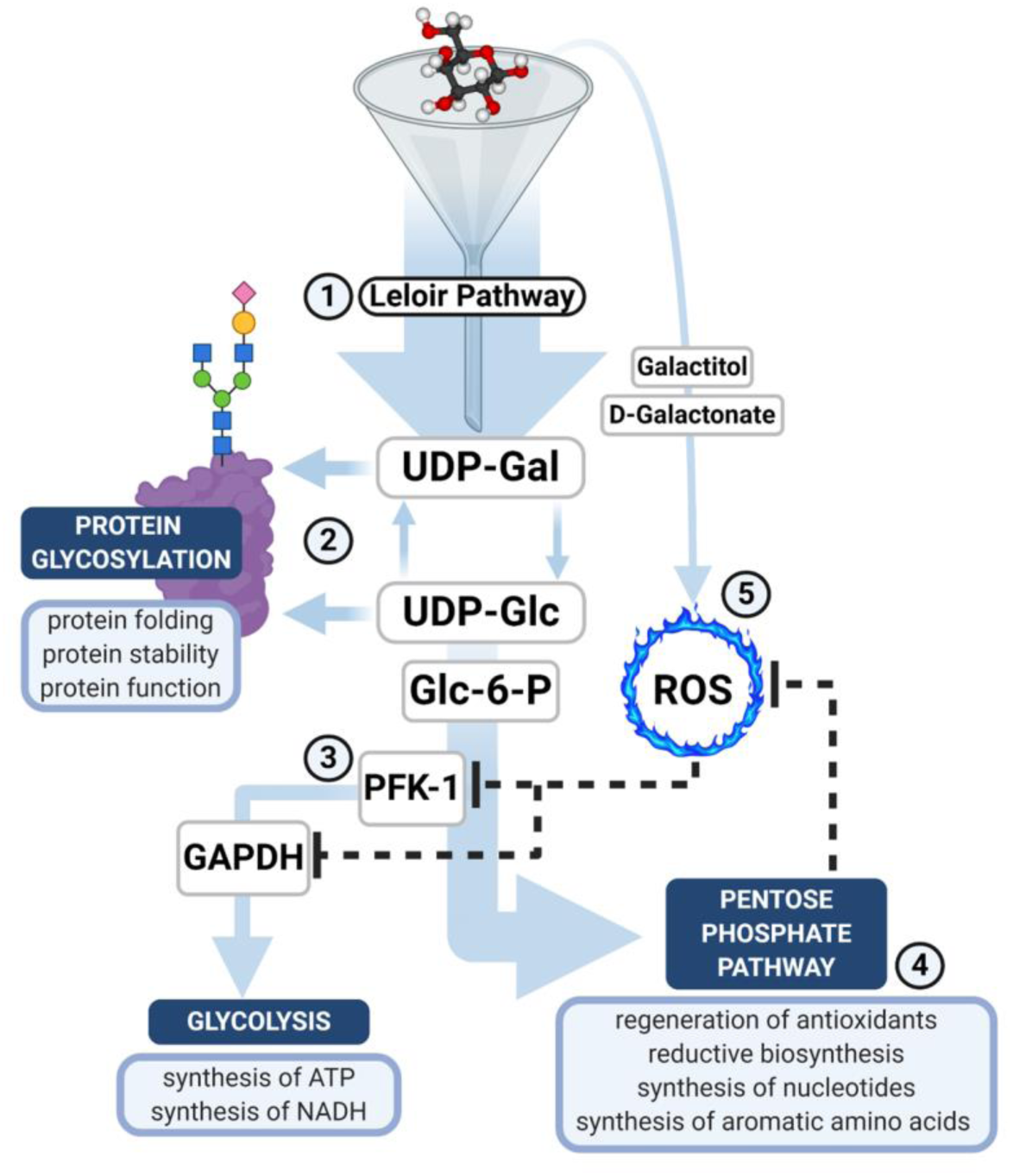
Schematic representation of potential biochemical pathways involved in galactose-induced hormesis. Leloir pathway **(1)** is considered as the most important metabolic route of galactose contributing biochemical substrates for **(2)** glycosylation pathways and the pool of cellular G-6-P that can either be routed towards **(3)** glycolysis to provide substrates for generation of ATP or towards **(4)** the pentose phosphate pathway to fuel regeneration of cellular antioxidants and reductive biosynthesis. Alternative metabolic pathways (e.g. metabolism to galactitol or D-galactonate) might act as regulators of the cellular galactose shuttle by stimulating the generation of ROS **(5)** that could then decide the fate of galactose in the cell by inhibiting critical regulatory enzymes such as PFK-1 and GAPDH, and hence rerouting galactose-generated G-6-P towards the PPP with the net effect being i) preserved glycosylation; ii) reduced metabolic activity and iii) regeneration of antioxidants, synthesis of nucleotides, and reductive biosynthesis.

## Discussion

### The response of systemic RRN indicators following single oral galactose treatment in rats

Analysis of the effects of a single dose of galactose administered by oral gavage on plasma and hippocampal RRN revealed several interesting patterns laying the groundworks for further investigation of mechanisms mediating both protective and harmful effects of galactose. Overall, the examination of plasma RRN suggests that acute increment of galactose availability is not sufficient to induce systemic oxidative stress following a single oral administration of 200 mg/kg. Interestingly, malondialdehyde (MDA), a single biomarker of oxidative stress that was significantly altered by the treatment indicates that acute oral galactose decreased rather than increased oxidative stress (**Fig 2C**). Malondialdehyde is an end-product produced by the degradation of arachidonic acid and polyunsaturated fatty acids often used as a biomarker of lipid peroxidation, a process of ROS-induced degradation of lipids. The increased concentration of MDA has been reported in various pathological processes such as Alzheimer’s disease, Parkinson’s disease, diabetes, cardiovascular diseases, cancer, and liver disease^58^. The biological background of this interesting phenomenon remains to be explained, however, our results indicate that oral galactose treatment can either diminish its production possibly by lowering lipid peroxidation, or potentiate the removal of this harmful electrophile aldehyde. One possible mechanism by which galactose could decrease the amount of MDA is by activating an aldehyde scavenger enzyme aldehyde dehydrogenase (**ALDH; EC 1.2.1.3**) known for its evolutionary conserved protective role in the cellular stress response^59^ either by hormetic stimulation through the generation of a moderate amount of lipid peroxides, or by replenishment of NADPH pool as further discussed in the text. The organ responsible for the observed effects remains to be determined, however, the liver may mediate some of the changes considering that hepatic tissue shows the highest initial capacity for galactose intake and metabolism^9^.

### Acute reorganization of hippocampal RRN following single oral galactose treatment

In contrast to systemic effects, RRN analysis of hippocampal tissue revealed a mild pro-oxidative shift with effect sizes and temporal patterns differing between individual redox subsystems. Paradoxically, a pro-oxidative shift was followed by a subsequent increment of the reductive potential in hippocampal tissue. For example, we observed a decrease of SOD activity after galactose treatment already 0.5h after the treatment (**Fig 3A**). A reduction in SOD activity has already been reported after chronic parenteral administration of galactose^16, 60, 61^, however, to our knowledge this is the first report of acutely decreased activity after oral administration in rats. Apart from SOD, the most pronounced change was observed for the H_2_O_2_ dissociation rate (**Fig 3B-C**). Chronic parenteral galactose treatment was reported to reduce catalase activity and various anti-aging therapeutics were demonstrated to counteract this change by increasing H_2_O_2_ dissociation capacity^18, 21, 61^. In contrast, we have found that acute oral treatment increased the H_2_O_2_ dissociation rate (**Fig 3B-C**). As ROS generation is considered to be the most important molecular mechanism driving galactose-induced senescence, we believe this paradoxical increment of H_2_O_2_ dissociation rate might be induced by substrate-mediated enzyme stimulation and suggests at least some of the changes responsible for the beneficial effects of galactose could be mediated by hormesis - a pathophysiological phenomenon describing a tendency of the biological system to respond to the stressor-induced homeostatic disbalance with protective overcompensation^62^. This is further supported by findings of increased peroxidase-like activity of cytochrome c (**Fig 3C**) as it has been demonstrated that its peroxidation capacity increases upon protein oxidation by reactive halogen species^63^. Hormetic principles have been extensively studied in the context of oxidative stress where continuous low-level ROS production has been identified as an important regulator of the cellular RRN capacity^64, 65^. For example, a mild oxidative stress-mediated protective effect has been observed for 3,5,4′-trihydroxy-trans-stilbene also known as resveratrol, a polyphenol with numerous health benefits found in red grapes^66^.

A hypothesis that a single oral administration can achieve hormetic levels of galactose in the hippocampus and stimulate redox overcompensation with a consequent increment of RRN capacity is further corroborated by the findings that the total amount of low molecular weight thiols remained practically unchanged by the treatment (**Fig 3D**), and that protein thiols and lipid peroxidation demonstrated a modest pro-oxidative shift fully compensated in the first 2 hours after galactose administration (**Fig 3E, Fig 3F**). Moreover, analyses of the overall redox balance measured by ORP and NRP suggest that galactose treatment resulted in an increased reductive capacity of the hippocampal environment despite a reduction in SOD activity (**Fig 3G-I**).

Spatiotemporal analysis of tissue reductive capacity by HistoNRP revealed that this hormetic effect was brain region-dependent as this short-term reductive hypercompensation was observed in ventral, but not dorsal hippocampus, and in temporal, but not in parietal cortical regions (**Fig 4A-C**). In this context, it would be interesting to explore the response of RRN subsystems in brain regions that demonstrated a linear dose-response reduction of antioxidant power following galactose treatment (**Fig 4C**) to see whether the lack of reductive shift was mediated by higher or lower galactose metabolic capacity with corresponding inverse ROS generation potential as this would further strengthen or denounce our hormetic hypothesis. Following the hypothesis that the observed effects were mediated by low levels of oxidative stress and adaptive response mechanisms, we further explored the NADP system as it has been suggested that NADPH-regulated disinhibition of the oxidative pentose phosphate pathway (PPP) flux serves as a rapid ROS detoxification metabolic system conserved across species^52, 67^. To combat oxidative stress cells upregulate the transcription of antioxidant defense system components^68^. However, before gene expression yields functional ROS scavengers, the survival of the cell depends on the short-term protective mechanisms that rapidly consume reducing equivalents mostly in the form of NADPH^52, 67, 69^. For this mechanism to work efficiently, cells had to evolve rapid and flexible biochemical rerouting strategies to supply substrates into the PPP responsible for the generation of NADPH^52, 53, 67, 69^. Analysis of hippocampal NADPH supports the hypothesis of galactose-induced mild oxidative stress with hormetic properties as the concentration of NADPH remained unchanged with a slight trend of the increment (**Fig 4D**). In the context of previously described RRN changes, we believe these findings indicate a potential increase in NADPH turnover with a metabolic rerouting-based replenishment of reducing equivalents used for the restoration of the antioxidant defense systems. This is further supported by the finding that total NADP dramatically increased following oral galactose treatment (**Fig 4E**) and that the ratio of NADPH and NADP decreased (**Fig 4F**).

Increased availability and turnover of NADPH observed here is in concordance with a classic cellular adaptive response to ROS since functional metabolic regulation providing substrates for NADP+ reduction not only fuels the most important cellular antioxidant systems such as glutathione peroxidase (**GPx; EC 1.11.1.9**) but also has the potential to reactivate catalase^70^. Furthermore, a dramatic increment of total NADP following galactose treatment is especially interesting. Oxidative stress has been reported to stimulate rapid phosphorylation of NAD+ to NADP+ by NAD+ kinase (**NADK; EC 2.7.1.23**) to increase substrate availability for the generation of NADPH. Nevertheless, this mechanism has mostly been described in plants and yeasts^71–73^. In contrast, it has been suggested that human cells generate NADPH to Fig ht oxidative stress not by increasing the NADP+ pool through NADK disinhibition, but through inducible NADP+-dependent dehydrogenase system-mediated replenishment of the reduced dinucleotide fraction^73–75^. This important difference is also reflected in the fact that yeast and plants have three well-defined isoforms of NADK^76^, while only one gene encoding cytosolic NADK has been identified in mammals in 2001^77^, and one additional mitochondrial variant has been described recently with yet undetermined pathophysiological functions^78^. The pathophysiological role of rodent NADK remains largely unexplored. The obvious importance of this enzyme in rodents is reflected in the fact that embryonic lethality arises as a consequence of loss of both NADK alleles in mouse^79^ and that spatiotemporal distribution of NADK activity plays an important role in the development of rat conceptus^80^. Apart from that, some indirect evidence exists that NADK might play an important role in oxidative stress response. Ethanol increases the synthesis of NADP+ in rat hepatocytes most likely as a result of increased NADK^81^ and butyric acid-induced oxidative stress increases NADP+ and the amount of blood cytosolic NADK and decreases NAD+^82^. Considering the biological relevance of NADP+ and NADK^73–75^, we believe our findings of dramatic increases of hippocampal NADP+ raise some important questions regarding the possible role of NADK on the crossroads of hippocampal RRN and metabolism. Following findings of mild increment of oxidative stress in rat hippocampus after a single oral galactose administration, compensated by metabolic shift with replenishment of NADP-derived reductive equivalents, we wanted to explore whether this metabolic shift would be reflected by a reduction of neuronal metabolic activity marker c-fos^83, 84^. This was based on the idea that rerouting of metabolic substrates towards the disinhibited oxidative branch of PPP in order to generate NADPH would reduce availability of metabolites for lower glycolysis, and ii) additional protective cellular regulatory mechanisms would be employed to further reduce the availability of NADH as it might have detrimental pathophysiological effects in the times of weakened reactivity of the cell due to its potential to generate ROS ^72^.

Our hypothesis was supported by findings of c-fos expression as acute oral galactose treatment reduced the expression in a time-dependent manner (**Fig 4G**). We also assumed that, apart from adaptive decrement of cellular activity, increased galactose availability would trigger expression of key enzymatic regulators of the Leloir pathway as this would allow the cell to balance energy production and NADPH replenishment in the environmental condition of prolonged galactose availability as G-6-P can be routed both towards lower glycolysis and PPP. As shown in **Fig 4H**, oral galactose treatment induced expression of GALK-1, an enzyme that catalyzes the first committed step of the Leloir pathway also in a time-dependent manner with increment being the most pronounced at the 2h time-point. Lastly, as Reutter’s group previously reported that one of the potential mechanisms by which galactose can exert its neuroprotective effects was by consumption of equivalents of ammonia through biosynthesis of amino acids from hexoses, we were interested to see how galactose-induced metabolic dynamics affected hippocampal glutamate homeostasis. The accumulation of glutamate has well-known neurotoxic effects and its importance has been discussed in the context of Alzheimer’s disease^85^. Nevertheless, glutamate also serves as one of the most important excitatory neurotransmitters and plays a major role in the regulation of neuronal energy by its engagement in the glutamate-glutamine cycle, a critical element of the astrocyte-neuron lactate shuttle, one of the prevailing contemporary theoretical frameworks of neuroenergetics^86^. Furthermore, glutamate metabolism is directly coupled to biochemical pathways responsible for the utilization of galactose as the first and rate-limiting enzyme of the hexosamine biosynthesis pathway (HBP), glutamine-fructose-6-phosphate transaminase (**GFAT; EC 2.6.1.16**) produces glutamate and glucosamine-6-phosphate from F-6-P and glutamine. It has been reported that the cellular pool of F-6-P increases in the conditions of increased galactose availability^4^ so it is possible that fueling the protein glycosylation pathways with galactose might also replenish cellular stores of glutamate through the process of transamination. This mechanism might be responsible for the restoration of glutamate stores that are acutely depleted after the treatment (**Fig 4I**) and corresponds to the increased expression of the Leloir pathway gatekeeper enzyme GALK1 (**Fig 4H**).

### A unique biochemical pattern mediates the protective effects of galactose in the conditions of limited substrate availability

Current evidence supports the concept that a unique galactose-induced metabolic pattern activates protective mechanisms in the conditions of limited substrate availability^4^. Sasaoka et al. demonstrated that galactose maintains mature protein glycosylation patterns more potently than glucose or mannose^4^. Considering the importance of protein glycosylation this finding indicates that galactose might preserve energy-sensitive cellular functions in times of reduced substrate availability. Maintenance of protein glycosylation patterns is especially important for growth factor signaling as both growth factors and their receptors are known to be heavily glycosylated and sensitive to perturbations of energy availability^87^. Interestingly, even the presence of trace amounts of galactose (0.3 mM) in the conditions of reduced substrate availability was able to correct starvation-induced disruption of insulin-like growth factor I (IGF-I) and epidermal growth factor (EGF) maturation and signaling in the HEK293 cells while the effect was not observed in the presence of the same amount of glucose^4^. The ability of galactose to maintain the function of growth factor signaling might even be directly related to its ability to stimulate rapid phosphorylation of NAD+ (**Fig 4E**) as it was recently reported that growth factors such as insulin or IGF-I exert direct control over NADK through Akt-mediated phosphorylation^88^. Furthermore, oxidative stress is a well-known consequence of reduced nutrient availability. Increased concentrations of superoxide radicals and H_2_O_2_ have been reported after starvation of glucose, glutamine, pyruvate, serum or amino acids, and starvation-induced generation of ROS has even been suggested as the main regulator of autophagy^89, 90^. In this context, the fact that galactose favors PPP over glycolysis makes it even more interesting for the preservation of cellular function during starvation, as besides growth factor signaling maintenance galactose provides substrates indispensable for the replenishment of cellular antioxidant defense reserves that are rapidly consumed by starvation-induced ROS. Diverting metabolism towards PPP might also activate other adaptive mechanisms for example by the rescue of glyceraldehyde-3-phosphate dehydrogenase (**GAPDH; EC 1.2.1.12**) through decreasing availability of its substrate glyceraldehyde-3-phosphate (G-3-P) as liberated GAPDH acts as a transcriptional regulator^56, 57^.

In summary, a large body of evidence supports the hypothesis that galactose has been selected by evolution as an ideal sugar to help the cells overcome fluctuations in the availability of energy substrates and maintain growth factors-related signaling. This hypothesis makes even more sense in the context of the fact that the importance of galactose seems to be the most pronounced in the brain, the most vulnerable and energy-dependent organ, and in the suckling period, the most vulnerable and energy-dependent period of life in mammals. In this context, it is possible that the same biochemical strategy that enables galactose to protect the brain from energy scarcity during the developmental period can be exploited to provide an alternative source of energy, replenish cellular stores of antioxidants, maintain protein glycosylation patterns, and rescue growth factor signaling during the process of neurodegeneration also characterized as a vulnerable period of reduced availability of energy substrates. This concept is supported by findings from our group as long-term oral galactose treatment both prevents^8^ and ameliorates ^38^ cognitive deficits in the rat model of sporadic Alzheimer’s disease induced by intracerebroventricular administration of streptozotocin (STZ-icv) characterized by disrupted insulin signaling in the brain^91^. Furthermore, oral galactose has been shown to normalize brain glucose metabolism both in STZ-icv ^38^ and in the Tg2576 model of Alzheimer’s disease ^42^.

### Galactose might exert its effects through hormesis: A hypothetical explanation of the relationship between protective effects of galactose and galactose-induced ROS

To conclude, ample evidence supports the hypothesis that galactose serves a unique biochemical role offering protection in times of limited substrate availability. However, evidence that continuous administration of galactose induces oxidative stress cannot be ignored, especially given the fact that a large number of groups exploit this effect to model aging in animals^92^. Based on the aforementioned, the inevitable question arises. How could it be that galactose induces oxidative stress, but also protects the cell from fluctuations in energy availability and increases the capacity of cells to tolerate ROS? We believe the findings reported here and the context provided by results reported by other groups provide a possible explanation for this perplexing paradox.

It could be that a specific metabolic reorganization following galactose administration is a direct consequence of the ability of galactose to induce tolerable amounts of cellular oxidative stress. In other words, it is possible that activation of specific biochemical machinery in the presence of galactose, which is responsible for the protective effects against oxidative stress and fluctuations of nutrient availability, is in fact hormetic in its nature. This hypothesis is supported by findings from other groups as metabolic dynamics following galactose treatment resemble biochemical reorganization following acute oxidative stress challenges. Disinhibition of the substrate flux into the oxidative branch of PPP with consequent replenishment of NADPH is considered to be an evolutionary conserved mechanism of protection against ROS present in different species ^52^ and the presence of galactose seems to favor the same substrate utilization pattern^56, 57^. Furthermore, oxidative stress-induced inhibition of glycolysis is further reinforced through the inactivation of gatekeeper enzymes, GAPDH and PFK-1^93^. In mammalian cells GAPDH serves as a pleiotropic sensor of oxidative stress through several mechanisms, the most important being direct ROS-induced intramolecular disulfide formation at active site cysteines and enzyme S-thiolation considered to be an important protective mechanism enabling the cell to rapidly resume glycolysis and ATP production after the oxidative stress subsides^93^. PFK-1, a critical glycolytic enzyme that serves a rate-limiting function and that is directly responsible for the first committed step of glycolysis is also inhibited following ROS challenge possibly through TP-53-induced glycolysis and apoptosis regulator (TIGAR)^93, 94^. Interestingly, the metabolic activity of GAPDH is decreased in the presence of galactose^56, 57^, and recent observations suggest that, unlike glucose that increases the activity of the glycolytic gatekeeper enzyme PFK-1, the presence of galactose affects neither concentration nor activity of the enzyme ^4^ enabling substrate shuttling towards the PPP. Furthermore, an increased expression of TIGAR has been reported in N2a cells when cultured in the presence of galactose^95^. Considering the aforementioned evidence, it is possible that the similarity of metabolic dynamics induced by galactose and ROS is not coincidental, and that the biochemical fate of galactose depends on its ability to activate the ROS-sensitive regulatory machinery of the cell. In this context, alternative galactose metabolic pathways, mainly catalyzed by aldose reductase (**AR; EC 1.1.1.21**) and galactose dehydrogenase (**EC 1.1.1.48**) are especially interesting as they are considered an important source of galactose-induced ROS. This is supported by findings reported by Li et al. who investigated the effect of galactose in malignant and non-malignant cell lines following the hypothesis that interruption of glycolysis might be an attractive approach to cancer therapy^95^. Interestingly, the effect of galactose in malignant cells was annihilated following pretreatment with sorbinil, a specific inhibitor of AR^95^, indicating that accessory metabolic pathways might play an important role in specific galactose-induced orchestration of biochemical machinery responsible for the observed effects. It also cannot be ruled out that galactose-induced hormesis might also be dependent on oxidative metabolism that generates a substantial amount of ROS as it has been shown that cells cultured in galactose media heavily rely on oxidative phosphorylation machinery, and this principle has been exploited for screening of fibroblasts from patients in whom respiratory chain-related defects have been suspected^96^ and for studying oxidative metabolism in human primary muscle cells^97^. In this context, galactose-induced protective effects might be related to mitohormesis ^98^. Further research more closely focused on supra acute effects of galactose, monitoring lactate-to-pyruvate ratios, and exploiting pharmacological and genetic manipulation with mitochondrial energetics might provide important information on whether and how mitohormesis might contribute to the overall observed hormetic effect.

### Further considerations and implications of the hormetic hypothesis of galactose in humans and animals

The hypothesis that the protective effects of galactose are inseparable from its stimulatory effect on ROS formation remains to be placed under scientific scrutiny. Nevertheless, if this hypothesis proves to be correct, its implications are likely important for understanding both physiological and pathophysiological functions of this unique monosaccharide. Better insights into how galactose orchestrates cellular metabolic response would help us understand the pathophysiological background in animal models of parenteral galactose-induced aging, protective effects of galactose in animal models of Alzheimer’s disease, dysregulation of biochemical pathways in case of inherited disorders of metabolism, and even its physiological importance. Regarding animal models, understanding the fine line between hormesis-mediated activation of protective cellular mechanisms, and detrimental consequences of the saturation of metabolic pathways would provide context important for understanding the perplexing results of animal experiments concerning galactose administration. Both the route of administration and the dose-response to galactose are critical components of galactose research that remain poorly understood. Because current evidence speaks in favor of galactose metabolic capacity of the cell as a critical factor for the determination of pathophysiological consequences of increased galactose availability, and that both factors exert significant influence over local galactose availability in different tissues, a proper understanding of these concepts is indispensable for interpretation of the results of animal studies on galactose. A plethora of studies have shown the detrimental effects of parenteral galactose^18, 92^, while oral galactose administration seems to exert beneficial effects at least in different animal models of Alzheimer’s disease^8, 38, 42^. Different explanations have been proposed considering these findings. For example, it has been shown that unlike intraperitoneal administration, oral galactose treatment increases the secretion of neuroprotective incretin glucagon-like peptide-1 (GLP-1)^38^ and it has been speculated that it might provide neuroprotective effects through modulation of gut microbiota and stimulation of afferent fibers of the vagus nerve^99^. Another possible explanation of these findings is a much larger and rapid increment of galactose availability following the intraperitoneal administration. For example, plasma galactose concentration is two times higher following intraperitoneal administration in comparison to orogastric gavage (208% vs 111% increment) in STZ-icv rats 15 minutes after the treatment^38^. In contrast, in this experiment, plasma galactose was unremarkable when compared to the untreated group even in the group sacrificed 30 minutes following treatment indicating most of the galactose is absorbed and/or metabolized in this time window (**Fig 2G**).

The importance of local dynamics of galactose availability is further corroborated by the findings that the beneficial effects of galactose on cognitive performance have also been reported following subcutaneous administration^39^. Furthermore, oral administration of galactose can also have detrimental effects in certain circumstances although the physiological gastrointestinal barrier acts to buffer the rapid entrance of large amounts of galactose into the body. In particular, it seems that long-term oral treatment has the potential to induce harmful changes when galactose is repeatedly given in bolus doses through orogastric gavage even at relatively low doses^40, 100^. From the hormetic hypothesis standpoint, local differences in cellular galactose availability that arise as a consequence of different experimental designs (often through pharmacokinetics) might explain the discrepant findings reported in the literature. For example, in our experiments, galactose achieved the best neuroprotective effect when it was dissolved in drinking water and administered throughout the day^8, 38^. Interestingly, plasma concentrations of galactose were either unchanged or even decreased in these animals in comparison to controls that didn’t drink galactose indicating that galactose metabolic capacity was not exceeded in these animals^38^. Another important caveat is to consider possible differences arising as a consequence of disturbed metabolism, rendering particular strains and animal models more susceptible or resistant to the observed hormetic effect. Although our preliminary results suggest neuroprotective properties of galactose are not unique for the STZ-icv model, and that they are also present in healthy animals [unpublished results], at the present moment most of the evidence is based on the findings from the rat model of sporadic Alzheimer’s disease with yet uncharacterized galactose metabolism. Furthermore, the concept of hormesis is intrinsically dependent on cellular compensatory capacity, and it is rational to anticipate some pathophysiological states will alter the efficacy of this approach by diminishing the ability of the cell to either withstand the original noxis or respond appropriately once the stimulus subsides.

### Conclusion

Based on the analysis of the effects of acute oral galactose treatment on plasma and hippocampal redox regulatory networks we propose that acute oral galactose treatment does not induce perturbations of the systemic redox networks in the dose of 200 mg/kg while the same dose is sufficient to induce a subtle pro-oxidative shift in the hippocampus that is (over)compensated in the first 2 hours following administration. The exact mechanism responsible for the observed (over)compensation remains to be further explored, however, rapid disinhibition of the oxidative PPP flux, a gradual reduction in neuronal activity, and increment of the expression of Leloir pathway gatekeeper enzyme GALK1 suggest a metabolic reorganization favoring the evolutionary conserved ROS detoxification metabolic system. Based on the observed changes, and in the context of findings reported by other groups, we propose a hormetic hypothesis of galactose action indicating that, at the biochemical level, protective effects of galactose might be inseparable from its ability to induce tolerable levels of cellular ROS.

### Limitations of our study

Being exploratory in its design, it is rational to conclude that our study did not achieve optimal power to detect potential alterations in all of the examined redox subsystems due to inherent limitations of particular biochemical methods or due to other reasons. For this reason, we clearly emphasize this throughout the text, and we discuss the observed changes without the “fixation on p-values”. Due to a relatively small number of animals and relatively large standard deviation, the trends we observed could just as well be meaningful and significant biological changes, and some significant differences could just as well be simple artifacts. We don’t consider this to be critical in this type of exploratory study where we were more focused on the association between particular subsystems and the effect on the redox regulatory network as a whole. Particular changes we observed remain in our focus and will be more clearly examined with the appropriate experimental design in our future experiments. Another critical limitation is the fact that we base our idea of the biochemical time-response on data obtained from individual animals randomized to different groups. In this regard, biochemical analysis of biological samples with repeated measurements of particular parameters would have been much more suitable for the assessment of acute galactose time response. Nevertheless, at this moment, it is impossible to conduct this kind of experiment. One option was a continuous sampling of extracellular fluid with the microdialysis probe placed in the particular hippocampal region, however, our experiments showed that i) placement of the probe, ii) pain associated with it, iii) flushing the tissue with artificial extracellular fluid, or iv) simply a biased recovery of particular molecules could affect the estimate of activity of the biological mechanisms we were interested in exploring in the dialysate (**Supplementary material**). The results obtained this way would therefore probably provide a poor representation of the biochemical changes that occur in reality. Another important limitation that should be emphasized is that no differentiation of neuronal and glial aspects was included in our study, and the results are only informative for the hippocampus in general. Differentiation of the hormetic effect on neuronal and glial cells remains the focus of our future experiments. Finally, we wish to draw the attention of the readers to one theoretical limitation related to the fact that for this purpose the term ROS was used throughout the text in a relatively general way to describe a large amount of pro-oxidative molecules involved in a similar and interrelated biochemical relationship with redox homeostatic systems. For some parameters we discuss, other terms would be more suitable (e.g. RCS – reactive carbonyl species)^101^.

## Author’s contributions

**JH, ABP, AK,** and **JOB** conducted the in vivo part of the experiment. **JH** established and validated biochemical methods for RRN analysis, and conducted measurements of SOD, LMWT, protein sulfhydryl content, TBARS, ORP, NRP, H_2_O_2_ dissociation rate, cytochrome c peroxidase-like activity, and HistoNRP. **JH, ABP** and **AK**, conducted measurements of NADP and NADPH, and Western blots for c-fos and GALK-1. **ABP, AK,** and **JOB** measured glutamate ELISA, glucose, and galactose. **JH** conducted data curation, analysis, and visualization, and wrote the manuscript. **ABP, AK, JOB, IK, DV, PR, and MSP** discussed the results, commented on the manuscript, and provided critical feedback. **MSP** (mentor of JH and PI of the lab) supervised the project and provided funding.

## Ethics committee approval

All experiments were conducted in concordance with the highest standard of animal welfare. Only certified personnel handled animals. Animal procedures were carried out at the University of Zagreb Medical School (Zagreb, Croatia) and complied with current institutional, national (The Animal Protection Act, NN135/2006; NN 47/2011), and international (Directive 2010/63/EU) guidelines governing the use of experimental animals. The experiments were approved by the national regulatory body responsible for issuing ethical approvals, the Croatian Ministry of Agriculture, and the Ethical Committee of the University of Zagreb School of Medicine.

