## Supplement 1 for "Is galactose a hormetic sugar? Evidence from rat hippocampal redox regulatory network"

### Supplementary Material

Materials and methods

#### Placement of the microdialysis probe

Placement of the microdialysis probe was performed on animals in deep anesthesia (ketamine 50 mg/kg / xylazine 5 mg/kg ip) 24 h prior to microdialysis to allow post-surgical recovery of the animals. In preparation of the probe placement, the insertion site on the animal's head was shaved, followed by a small incision of the skin and drilling of the skull for insertion of the cannula and to secure the probe in place using 3 screws. A small catheter containing a semipermeable membrane was placed stereotaxically into the hippocampus. The probe was further sealed to the animal's head using GC FujiCEM2 dental cement (GC Corporation, Japan). The microdialysis probe was perfused internally with artificial cerebrospinal fluid at a fixed rate of 2 µl x $min^{-1}$ using the microdialysis pump by CMA (Harvard Bioscience, USA). After equilibration between the two compartments, the composition of the collected microdialysate reflects that of the brain extracellular fluid. Metamizole (Dipyrone) [100 mg/kg] was used for postoperative analgesia in concordance with the guidelines for pain management in laboratory animals [1]. Contralateral hippocampus was used for the analyses. All analyses were conducted as described in the Methods section in the main text.

Preliminary analysis of the redox regulatory network (RRN) in plasma and hippocampus of rats with microdialysis probe demonstrated that i) brain trauma induced by placement of the probe; ii) anesthesia; iii) pain induced by the surgery; iv) postoperative analgesia, or all of the above could affect at least some of the parameters of the RRN. Microdialysis (MD)-treated animals had increased reductive capacity and low molecular weight thiol (LMWT) concentration in plasma, and increased concentration of thiobarbituric acid reactive substanes (TBARS), a common lipid peroxidation biomarker (**Fig S1**). RRN changes in hippocampal homogenates were less pronounced, however most of the measured parameters demonstrated increased dispersion (**Fig S2**). Change in plasma total antioxidant capacity, LMWT and lipid peroxidation have all been discussed in the literature in the context of the pathophysiological response to traumatic brain injury [2]. Furthermore, dipyrone has been recognized as a potent antioxidant and scavenger of reactive nitrogen species (RNS)[3], and it has been proposed that ketamine is able to rapidly decrease reactive oxygen species (ROS) production [4]. On the other hand, ketamine-xylazine anaesthesia has been recognized as „oxidative anesthesia" as it can induce lipid peroxidation, decrease activity of catalase and superoxide dismutase, and increase the activity of glutathione peroxidase [5]. In this regard, and in the context of the findings presented below, we decided that microdialysis was not the best approach considering operation, tissue perfusion, and pharmacological agents that have to be used in order to successfully install and use the probe have all been recognized as modulators of redox homeostasis.

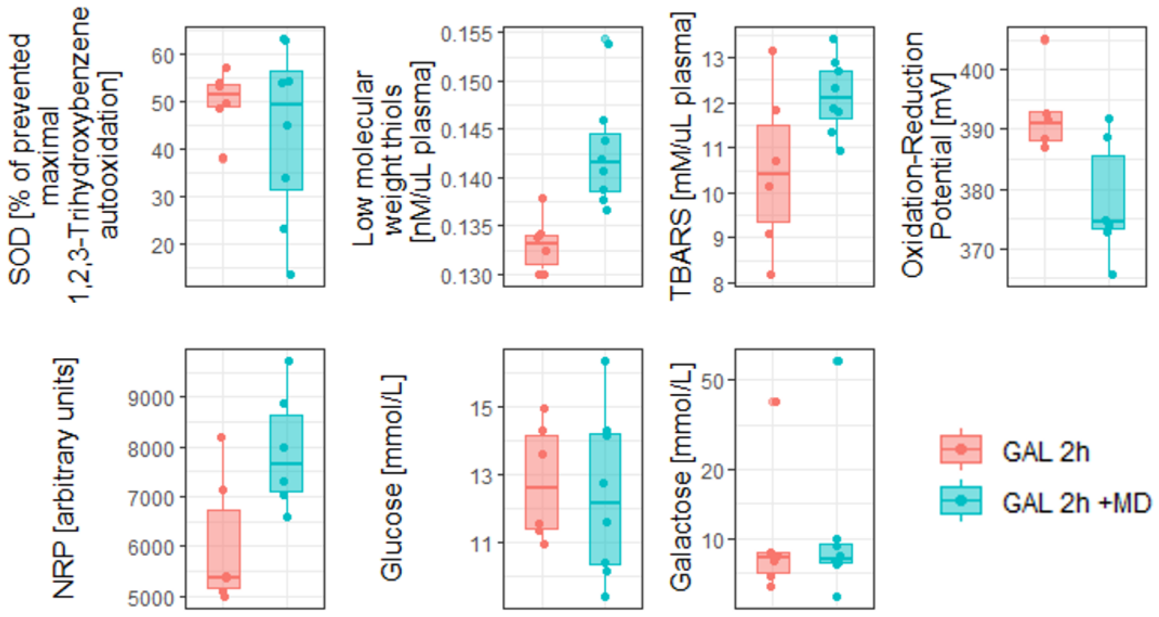


**Fig S1. Comparison of the activity of constituents of the plasma redox regulatory network, concentration of glucose and galactose in rats 2h after 200mg/kg orogastric galactose treatment with or without microdialysis probe placed in hippocampal tissue 24 hour before the experiment**.

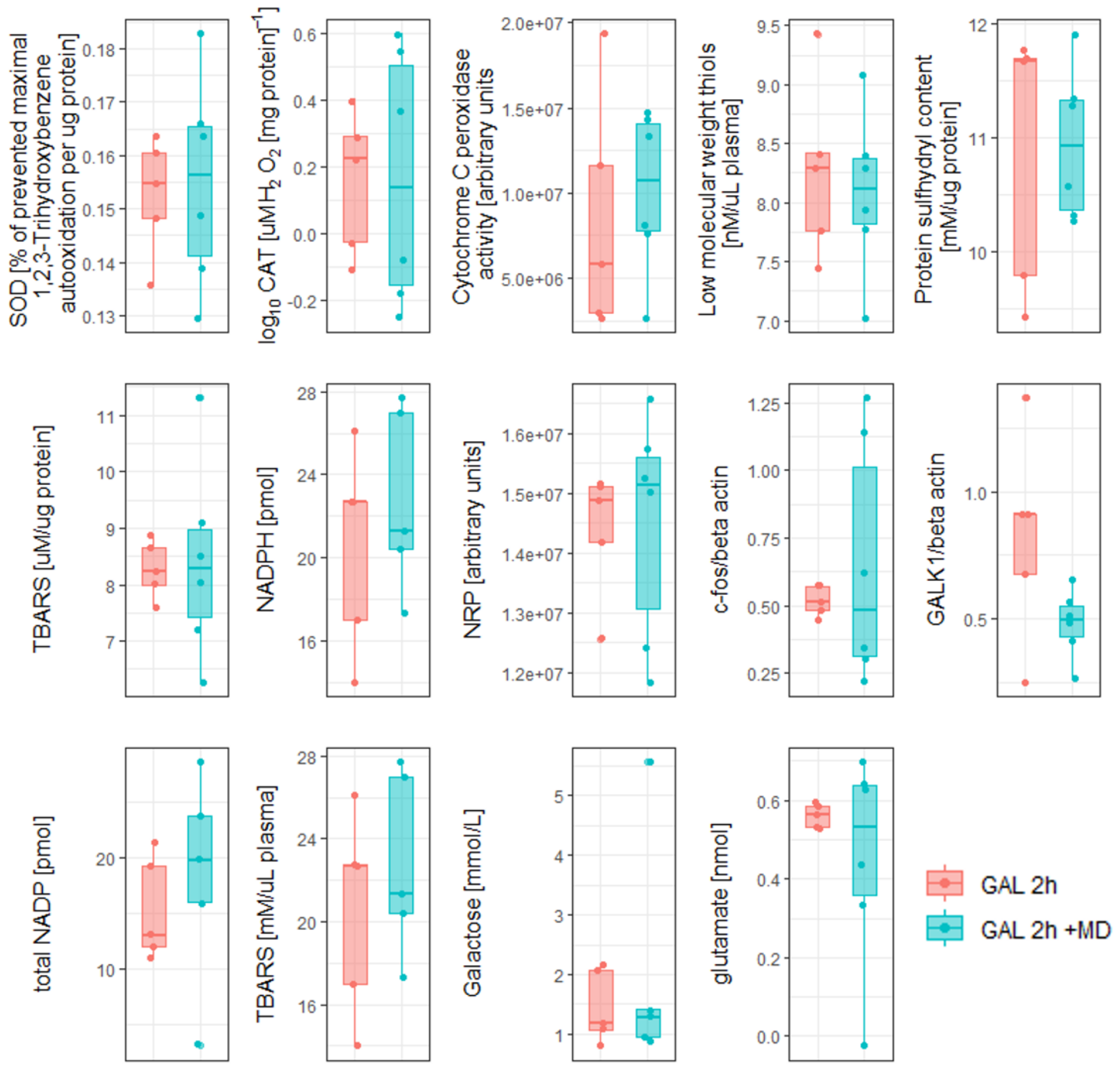


**Fig S2. Comparison of the activity of constituents of hippocampal redox regulatory network, in rats 2h after 200mg/kg orogastric galactose treatment with or without microdialysis probe placed in hippocampal tissue 24 hours before the experiment.**

Homolak et al. **Is galactose a hormetic sugar? Evidence from rat hippocampal redox regulatory network (2021)**
